# Kaptive 2.0: updated capsule and LPS locus typing for the *Klebsiella pneumoniae* species complex

**DOI:** 10.1101/2021.11.05.467534

**Authors:** Margaret M. C. Lam, Ryan R. Wick, Louise M. Judd, Kathryn E. Holt, Kelly L. Wyres

## Abstract

The outer polysaccharide capsule and lipopolysaccharide antigens are key targets for novel control strategies targeting *Klebsiella pneumoniae* and related taxa from the *K. pneumoniae* species complex (KpSC), including vaccines, phage and monoclonal antibody therapies. Given the importance and growing interest in these highly diverse surface antigens, we had previously developed Kaptive, a tool for rapidly identifying and typing capsule (K) and outer lipopolysaccharide (O) loci from whole genome sequence data. Here, we report two significant updates, now freely available in Kaptive 2.0 (github.com/katholt/kaptive); i) the addition of 16 novel K locus sequences to the K locus reference database following an extensive search of >17,000 KpSC genomes; and ii) enhanced O locus typing to enable prediction of the clinically relevant O2 antigen (sub)types, for which the genetic determinants have been recently described. We applied Kaptive 2.0 to a curated dataset of >12,000 public KpSC genomes to explore for the first time the distribution of predicted O (sub)types across species, sampling niches and clones, which highlighted key differences in the distributions that warrant further investigation. As the uptake of genomic surveillance approaches continues to expand globally, the application of Kaptive 2.0 will generate novel insights essential for the design of effective KpSC control strategies.

**Significance as a BioResource to the community:** *Klebsiella pneumoniae* is a major cause of bacterial healthcare associated infections globally, with increasing rates of antimicrobial resistance, including strains with resistance to the drugs of last resort. The latter have therefore been flagged as priority pathogens for the development of novel control strategies.

*K. pneumoniae* produce two key surface antigen sugars (capsular polysaccharide and lipopolysaccharide (LPS)) that are immunogenic and targets for novel controls such as a vaccines and phage therapy. However, there is substantial antigenic diversity in the population and relatively little is understood about the distribution of antigen types geographically and among strains causing different types of infections. Whereas laboratory-based antigen typing is difficult and rarely performed, information about the relevant synthesis loci can be readily extracted from whole genome sequence data. We have previously developed Kaptive, a freely available tool for rapid typing of *Klebsiella* capsule and LPS loci from genome sequences.

Kaptive is now used widely in the global research community and has facilitated new insights into *Klebsiella* capsule and LPS diversity. Here we present an update to Kaptive facilitating i) the identification of 16 additional novel capsule loci, and ii) the prediction of immunologically relevant LPS O2 antigen subtypes. These updates will enable enhanced sero-epidemiological surveillance for *K. pneumoniae,* to inform the design of vaccines and other novel *Klebsiella* control strategies.

**Data summary:**

1. The updated code and reference databases for Kaptive are available at https://github.com/katholt/Kaptive
2. Genome accessions from which reference sequences of novel K loci were defined are listed in Supplementary Table 1, and genomes from which these loci were detected (along with the corresponding Kaptive output) are listed in Supplementary Table 2.
3. Accessions for the genomes screened for O types/subtypes (along with the corresponding Kaptive output) are listed in Supplementary Table 3.

**The authors confirm all supporting data, code and protocols have been provided within the article or through supplementary data files**.

**Repositories:** *Repositories:* Genome sequence from which the novel K locus KL182 was defined has been deposited under the accession JAJHNT000000000.

## Introduction

The *Klebsiella pneumoniae* species complex (KpSC) is a group of closely related Gram-negative bacterial taxa including the opportunistic pathogen, *Klebsiella pneumoniae* [1]. The ‘K’ in the ESKAPE pathogens, *K. pneumoniae* is considered one of the six most important causes of drug resistant healthcare-associated infections [2], and antimicrobial resistant strains contribute significantly to the total burden of communicable disease in high income countries [3]. In low- and middle- income countries *K. pneumoniae* is also recognised as the leading cause of Gram-negative neonatal sepsis, contributing to 10% of total neonatal sepsis deaths [4, 5]. *K. pneumoniae* with resistance to the third-generation cephalosporins and carbapenems are disseminating globally and of particular concern because they cause infections with very limited treatment options. As a consequence, there is increasing interest in developing novel anti-KpSC control strategies such as vaccines, phage and monoclonal antibody therapies [6–9].

The KpSC polysaccharide capsule and lipopolysaccharide (LPS) antigens are pathogenicity factors [10–13] and among the key targets for novel control strategies [7,14–18]. Of particular interest, there is mounting evidence that capsule and/or LPS immunisation can elicit a protective immune response, and several anti-KpSC capsule and/or LPS vaccines have entered clinical trials [19–23]. However, considerable antigenic diversity exists within the KpSC population (>77 serologically defined capsule types [24–26], >8 LPS O antigen serotypes [16, 27], and many more predicted on the basis of genomic data [28, 29]), and there is a paucity of sero-epidemiological knowledge, which hampers efficient vaccine design. While traditional KpSC serological typing techniques are technically challenging and rarely performed, information about capsule and O antigen serotypes can be rapidly extracted from whole genome sequence (WGS) data by typing the corresponding capsule (K) and O biosynthesis loci [29, 30].

The K locus comprises a ~10-30 kbp region of the chromosome and ~10-30 genes. The conserved capsule synthesis and export machinery genes are located in the 5’-(*galF, cpsACP, wzi, wza, wzb, wzc*) and 3’-(*ugd*) most regions, and flank the genes encoding the capsule-sugar-specific synthesis machinery, Wzx capsule-specific flippase, and Wzy repeat unit polymerase [31, 32], The O locus comprises a ~7-13 kbp region of the chromosome between the K locus and the *hisE* gene [28]. All O loci identified to-date contain the *wzm* and *wzt* genes encoding the membrane transporter complex [33], but the relative positions of these genes is not consistent [28].

Comparison of the K loci of the 77 original capsule serotype reference strains showed that all but two could be distinguished by a unique combination of genes in the centre of the locus, i.e. there was an approximate one-to-one relationship between K locus and serologically-defined K type [32]. In light of these findings, we previously leveraged a collection of >2500 KpSC genomes to identify additional K loci, defined on the basis of unique gene content [28, 29]. A total of 59 loci were identified, and labelled KL101-KL159. While the corresponding serotypes remain uncharacterised, we predict that these loci encode distinct capsule types that likely correspond to the majority of the 10-70% of strains that have been deemed non-typeable and/or cross-reactive via serological typing techniques [34–36].

To facilitate the rapid identification of K loci from KpSC whole genome sequences, we developed a tool known as Kaptive [29], which uses a combination of BLASTn and tBLASTn search to identify the best matching locus from the reference database, and provide an indication of the match confidence. We recommend reporting matches of ‘Good’ confidence or higher. Low or no confidence matches generally result from sequencing and/or assembly problems that cause the K locus to be split across multiple assembled contigs, but can also represent novel loci that were not captured in the original discovery genome set. Indeed, subsequent studies have identified 11 additional loci that have been incorporated into the database ([37] and see details at github.com/katholt/kaptive), resulting in a total of 146 loci defined to-date. We anticipate the discovery of additional novel loci, particularly given the availability of greater numbers of KpSC genomes from diverse sources and locations. Notably, our recent analysis of >13,000 publicly available KpSC genomes identified 19% with low or no confidence K locus matches that remain to be explored [38].

The majority of distinct LPS O antigens are also associated with distinct O loci (also known as *rfb* loci), distinguished on the basis of gene content [39–42], However, key exceptions are strains expressing the O1 and O2 antigens, both of which can be associated with either of two distinct O loci, O1/O2v1 and O1/O2v2 [15, 41, 43]. Expression of either locus results in the production of an O2 antigen (D-galactan I or III), that can be converted to O1 by addition of a D-galactan II repeat unit by WbbYZ (encoded elsewhere in the genome) [44].

In 2018 we extended Kaptive to support KpSC O locus typing and included discrimination of the O1 and O2 types by tBLASTn search for the *wbbYZ* genes, reported as shown in Table 1 [30]. However, recent experimental evidence has shown that only the WbbY protein is needed to convert O2 to O1 [45]. Additionally, the genetic determinants of four distinct O2 antigen subtypes have now been fully elucidated (O2a, O2afg, O2ac, O2aeh) [45, 46]. The O2a antigen comprises alternating repeat units of α-(1-3)–linked galactopyranose (Gal*p*) and β-(1-3)–linked galactofuranose (Gal*f*) residues (D-galactan I) [40]. O2afg comprises O2a with a (1-4)-linked Gal*p* side chain, while O2aeh comprises O2a with a (1-2)-linked Gal*p* side chain [43]. Like O1, O2ac comprises O2a or O2afg with an additional repeat unit covalently linked to the non-reducing terminus (it also seems likely that O2aeh can be modified in this way). While the O1 repeat unit comprises [-3)-α-D-Gal*p*-(1-3)-β-D-Gal*p*-(1-] (also known as D-galactan II) [40], the O2ac repeat unit comprises [-3)-β-D-GlcpNAc-(1-5)-β-D-Gal*f*(1-] [43, 46]. These subtypes are associated with specific combinations of O loci (O1/O2v1, O1/O2v2 and a novel locus identified in the O2aeh type strain, here called O1/O2v3) with/without additional genes located elsewhere in the genome (*wbbY, gmlABD* and *wbmVW* [45, 46], see **Table 1**). There is emerging evidence that these subtypes differ in terms of immunogenicity [15, 17, 47] but little is known about their distribution in the KpSC population, which may have implications for the design of vaccines or monoclonal antibody therapies targeting KpSC LPS.

**Table 1:** Genetic determinants of *Klebsiella pneumoniae* species complex O1 and O2 outer lipopolysaccharide antigens as reported in Kaptive.

| O locus | Extra genes | Kaptive <v2.0 | Kaptive ≥v2.0 | |
| --- | --- | --- | --- | --- |
|  |  | Locus<sup>a</sup> | Locus<sup>a</sup> | Type<sup>b</sup> |
| O1/O2v1 | none | O2v1 | O1/O2v1 | O2a |
| O1/O2v2 | none | O2v2 | O1/O2v2 | O2afg |
| O1/O2v3 | none | N/A | O1/O2v3 | O2a |
| O1/O2v1 | *wbbYZ*<sup>c</sup> | O1v1 | O1/O2v1 | O1 |
| O1/O2v2 | *wbbYZ*<sup>c</sup> | O1v2 | O1/O2v2 | O1 |
| O1/O2v3 | *wbbYZ*<sup>c</sup> | N/A | O1/O2v3 | O1 |
| O1/O2v1 | *wbbY* OR *wbbZ* | O1/O2v1 | N/A | N/A |
| O1/O2v2 | *wbbY* OR *wbbZ* | O1/O2v2 | N/A | N/A |
| O1/O2v3 | *wbbY* OR *wbbZ* | N/A | N/A | N/A |
| O1/O2v1 | *wbmVW* | N/A | O1/O2v1 | O2ac |
| O1/O2v2 | *wbmVW* | N/A | O1/O2v2 | O2ac |
| O1/O2v3 | *wbmVW* | N/A | O1/O2v3 | O2ac |
| O1/O2v1 | *gmlABD* | N/A | O1/O2v1 | O2aeh |
| O1/O2v2 | *gmlABD* | N/A | O1/O2v2 | O2aeh |
| O1/O2v3 | *gmlABD* | N/A | O1/O2v3 | O2aeh |
| O1/O2v1 | *wbbY* AND *wbmVW* | N/A | O1/O2v1 | O1 (O2ac)<sup>d</sup> |
| O1/O2v2 | *wbbY* AND *wbmVW* | N/A | O1/O2v2 | O1 (O2ac)<sup>d</sup> |
| O1/O2v3 | *wbbY* AND *wbmVW* | N/A | O1/O2v3 | O1 (O2ac)<sup>d</sup> |

<sup>a</sup> as reported in the 'Best match locus' column in the Kaptive output
<sup>b</sup> predicted antigenic serotype reported in the 'Best match type' column in the Kaptive output (v2.0 and above)
<sup>c</sup> Kaptive v2.0 and above check only for *wbbY*.
<sup>d</sup> predicted antigenic serotype likely O1 but may also be O2ac (there is currently no corresponding type strain with *wbbY* and *wbmVW*)
N/A - not applicable

Here we report; i) an update to the KpSC K locus database to include 16 novel loci identified from a high-throughput screen of >17,000 publicly available genomes; and ii) an updated version of Kaptive that can rapidly distinguish O2 subtypes to support enhanced LPS sero-epidemiological investigations. We apply this update to explore the distribution of O (sub)types between KpSC species and among isolates from different sources and clonal groups (CGs), including the well-known globally distributed multi-drug resistant clones and the hypervirulent clones associated with severe community-acquired disease.

## Methods

### Identification of novel K loci

*We* have previously reported genotyping information for 13,156 publicly available KpSC genomes (including species, multi-locus sequence types (STs), K loci and K locus confidence calls) [38]. Here we leverage these data in combination with 3,958 additional published KpSC genomes and 638 from our private collection (**Table 2**) for which the corresponding sequence reads were *de novo* assembled using Unicycler vO.4.7. Species and STs were determined using Kleborate v2.0.3 [38] and K loci identified using Kaptive v0.7.3 [29].

**Table 2:** Sources of *K. pneumoniae* species complex (KpSC) genomes from which novel K loci were identified.

| Collection description | Reference | N Genomes (N low confidence K loci<sup>a</sup>) |
| --- | --- | --- |
| Curated, publicly available KpSC isolate genomes | [38] (also see refs therein) | 13156 (1802) |
| KpSC isolate genomes from diverse niches in a single city in Italy | [48] | 1965 (75) |
| Blood and urinary tract KpSC isolate genomes from Norwegian hospitals | [49] | 868 (358) |
| Clinical KpSC isolate genomes from an Australian hospital | This study | 638 (18) |
| Community rectal carriage KpSC isolate genomes from Norway | [50] | 484 (36) |
| KpSC neonatal sepsis isolate genomes from south east Asia and Africa | [5] | 276 (93) |
| KpSC isolate genomes from pigs and pig farmers in Thailand | [51] | 253 (31) |
| *Klebsiella variicola* from various sources | [52] | 112 (6) |
| **Total** |  | **17752 (2419)** |

<sup>a</sup> Genomes harbouring K loci for which the Kaptive confidence call was 'Low' or 'None', and subsequently included in this study.

Genomes for which the Kaptive K locus confidence call was “Low” or “None” were included for further analysis as follows: genomes for which the Kaptive output did not indicate a fragmented K locus assembly (i.e. the K locus problems column did not contain ‘?’) were subjected to manual inspection using Bandage v0.8.1 [53] to visualise the BLASTn coverage to the best match K locus and assess if the genome truly harboured the best match locus, a variant thereof (insertion sequence (IS) or deletion variant) or a putative novel locus. Genomic regions corresponding to putative novel loci were extracted and clustered using CD-HIT-EST v4.8.1 (default parameters) [54]. A single representative of each cluster was; a) annotated using Prokka v1.14.6 [55] with a reference database of known KpSC K locus genes; b) subjected to BLASTn search for known KpSC K locus genes and those annotated in each of the other putative novel K loci. Inspection of the BLASTn results highlighted putative novel loci with similarity to each other and/or existing loci (i.e. those with BLASTn hits ≥80% identity and ≥80% coverage to multiple capsule-sugar-specific synthesis genes from the same reference locus). The corresponding loci were subsequently compared by BLASTn and visualised with the Artemis Comparison Tool v18.0.2 [56] to clarify if they were distinct or should be considered as IS or deletion variants of the same locus.

Annotations were manually curated for one representative of each of the final set of distinct novel loci (**Supplementary Table 1**). Where possible, sequences without IS transposase annotations were preferentially selected for curation, as is recommended in order to prevent Kaptive reporting spurious gene matches to transposases that may be present in multiple copies in any location of a query genome. Where no IS transposase-free representative was available (n=5 loci, subsequently assigned KL173, KL176, KL181, KL184 and KL185), full length ISs were identified by BLASTn search of the ISfinder database [57] and manually deleted along with their associated direct repeats and one copy of the associated target site duplicated repeat to obtain a putative IS-free reference sequence. Curated locus annotations were added to the reference database and Kaptive was rerun on all genomes to determine the prevalence of novel loci (**Supplementary Table 2**). Only Kaptive locus calls with confidence ‘Good’ or better are reported. Visual comparisons of the novel K loci and annotated coding sequences (CDS) were generated with clinker v0.0.21 [58].

### Implementation of O2 subtyping

Kaptive takes as input a query genome assembly and a reference locus database in GenBank format. Loci are identified by a note field within the sequence source information in the format:

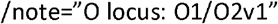

Additional genes relevant to KpSC O typing (i.e those located outside of the O locus) are marked as follows:

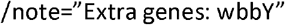

In Kaptive 2.0 we have implemented an additional note field for reference loci that indicates the corresponding serotype. E.g. for the O5 locus the corresponding O type is known to be O5 and hence the corresponding note fields read:

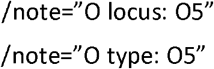

For loci identified from genome data and defined on the basis of gene content alone, e.g. OL1O1, there is no known serotype and hence the corresponding note fields read:

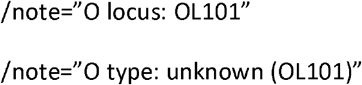

For the O1 and O2 loci, the corresponding serotypes/subtypes are distinguished by the presence/absence of additional genes located elsewhere in the genome, as is now indicated by the use of the term ‘special logic’ in the serotype notes field e.g:

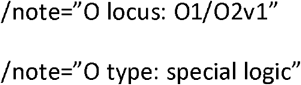

After identifying the best match locus from the database (the locus with the highest BLASTn coverage), Kaptive 2.0 will extract the corresponding type information from the GenBank annotation. When the type is indicated as ‘special logic’ Kaptive will perform tBLASTn search for all coding sequences within database entries marked with the note field ‘Extra genes’ (*wbbY*, Genbank accession MG458672.1; *wbmVW*, accession MG602074.1; *gmlABD*, accession MG458670.1). tBLASTn hits exceeding the minimum identity and coverage cut-offs (default 80% and 90%, respectively) are interpreted to indicate the presence of the corresponding gene(s). The reported O type is determined by the combination of best match O locus and any extra gene(s) detected, as specified in the “Klebsiella_o_locus_primary_reference.logic” file and shown in **Table 1**. In the event that a combination is not represented in the logic file, Kaptive will report the O type as “unknown.”

### Updated locus and serotype reporting

In the previous version of Kaptive the distinction between the KpSC O1 and O2 types was indicated in the output table in the ‘Best match locus’ as shown in **Table 1**. However, we have noted that this has resulted in confusion in the community regarding the distinction between O loci and O types. In order to clarify these differences and ensure that all relevant information is included in the output, Kaptive 2.0 reports the best match locus and the predicted serotype in separate columns in the output (see **Table 1**). The three distinct O loci associated with the O1 and O2 (sub)types are reported simply as O1/O2v1, O1/O2v2 and Ol/O2v3 in the ‘Best match locus’ column, meaning that genomes harbouring the same O locus can be easily identified even when predicted to express a different O type due to variation elsewhere in the genome.

The same approach for distinguishing ‘Best match locus’ from ‘Best match type’ is applied to all databases parsed by Kaptive 2.0 in order to aid interpretation of the output and clarify what is known about the relationships between loci and serotypes. In particular we hope this will clarify where serotypes are or are not known. To enable backwards compatibility for reference locus database that do not contain serotype note fields, the corresponding types are left blank if no type information is available.

### Exploring the distribution of O (sub)types

***We*** used Kaptive 2.0 to identify O loci and O types for 10,734 non-redundant KpSC genomes in the curated collection reported previously [38]. Additionally, we supplemented this collection with two systematically collected datasets of KpSC from underrepresented sources: 484 human gut carriage isolate genomes reported by Raffelsburger *et al* [50], and 875 non-human associated isolate genomes reported by Thorpe *et al* [48]. (We excluded human-associated KpSC genomes reported as part of the latter study because these data were biased by a significant overrepresentation of the ST3O7 and ST512 clones that were circulating in the local hospital and were already represented in high numbers in our curated genome collection.) Only Kaptive calls with confidence ‘Good’ or better are reported.

## Results and Discussion

### Identification and characterisation of 16 novel K loci

A total of 2,419 of 17,752 KpSC genomes (13.6%) had low or no confidence Kaptive K locus calls (**Figure 1**), of which the vast majority (2,129; 88%) were indicated as fragmented and 37 were excluded because the K loci were likely to harbour an overrepresentation of homopolymer repeat errors due to the assembly approach comprising Oxford Nanopore Technologies read data only. Following manual inspection, 148 of the 253 remaining genomes were considered to match to known K loci and 45 were unresolvable because the K locus was fragmented across multiple assembly contigs. The former had not been confidently typed by Kaptive for a variety of reasons including IS insertions, small- or large-scale deletions and/or numerous frameshift mutations that were interpreted as missing genes. After exclusion of these genomes, 60 putative novel loci remained and were grouped into 22 sequence clusters at 90% nucleotide identity. Sixteen clusters were determined to represent true distinct and novel K loci, while 6 were identified as additional IS/deletion variants of other putative novel loci or existing loci (**Figure 1**, see **Methods**).

**Figure 1:**
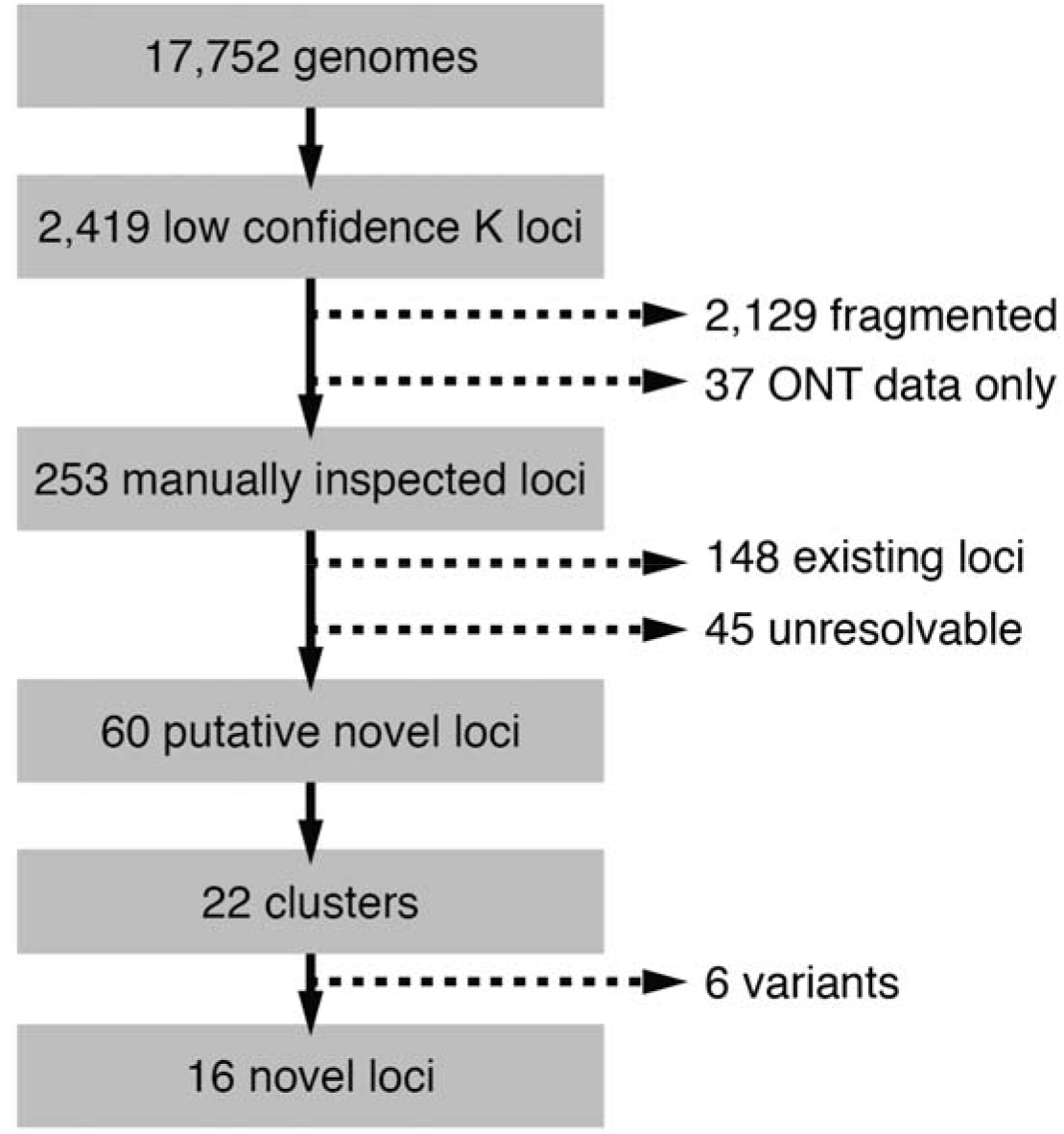
Process of identification of novel K loci from 17,752 *K. pneumoniae* species complex genomes. Candidate genomes were identified as those with Kaptive K locus confidence calls ‘Low’ or ‘None,’ and were iteratively filtered to remove; i) fragmented or low quality locus sequences including 37 assembled using only Oxford Nanopore Technologies (ONT) data; ii) true matches to existing K loci; iii) insertion sequence and/or deletion variants of existing loci or other putative novel loci (see **Methods** for full details).

The final set of 16 novel loci were assigned KL171-KL186, and ranged in length from 22,624 to 30,215 bp and contained 19 to 25 genes (**Figure 2**). All loci contained the conserved capsule synthesis and export machinery genes; *galF, cpsACP, wza, wzb, wzc, gnd* and *ugd*; however, only 14 loci harboured *wzi* (present in >98% of all previously reported KpSC K loci [29]). The Wzi protein is not considered essential for capsule production but plays a role in capsule surface attachment such that mutants lacking *wzi* produce lower levels of bound and higher levels of cell-free polysaccharide [59]. In the novel KL175 and KL176 loci, the *wzi* gene appeared to be replaced by four coding sequences predicted to encode two hypothetical proteins, a GfcC protein and a putative lipoprotein YjbH precursor. N=3/4 of these sequences had 76-84% sequence identity to those present in the KL33 and KL4O reference loci, wherein *wzi* is also absent [32].

**Figure 2:**
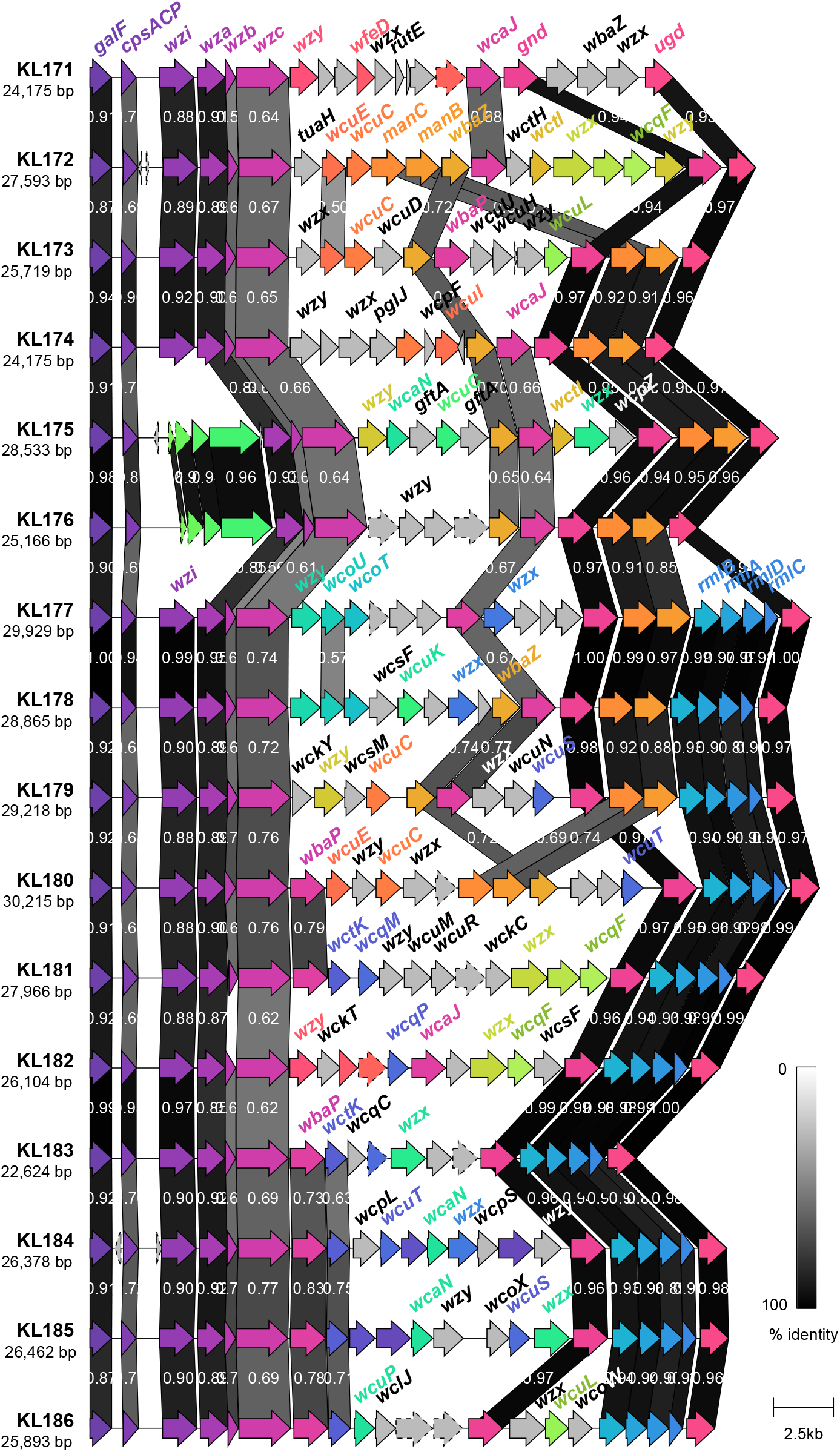
Genetic structure of novel K loci identified in this study. Coding sequences are represented by arrows, labelled by their gene name where applicable, and coloured by homology (sequence identity ≥30%). Coding sequences predicted to encode hypothetical proteins are represented by arrows with a dashed outline. Shading between the K loci represents regions of similarity between coding sequences as identified by clinker [58], and the level of similarity indicated in the legend.

GfcC proteins are known to be involved in the export of *Escherichia coli* group 4 capsules where they form a pore that allows the capsule to cross the cell membrane in a role similar to that of the KpSC Wza protein [60]. However, apparently intact copies of *wza* were also present in each of the KL33, KL40, KL175 and KL176 loci, so it is unclear what if any role GfcC plays in the export of these capsules. Similarly, the role of the YjbH precursor protein is unclear. YjbH is indicated as a regulator of exopolysaccharide expression in *E. coli* but it is encoded as part of the *yjbEFGH* [61] operon that was absent from these KpSC K loci.

As expected, all novel loci harboured a variant of *wzx* and of *wzy*, encoding the capsule flippase and repeat unit polymerase, respectively, as well as either *wcaJ* or *wbaP* (encoding the initiating glycosyl transferases) [31]. Ten loci (63%) harboured the *rmlBADC* capsule-specific sugar synthesis genes responsible for the inclusion of dTDP-L-rhamnose in the capsule and 9 (57%) harboured *manBC* associated with the inclusion of GDP-D-mannose [32]. The remaining capsule-specific sugar synthesis genes were each found in ≤5 loci each.

We assessed the prevalence of the novel loci among the public genome collections (**Table 2**). Most (n=11; 69%) were rare, identified in ≤5 genomes. However, KL174, KL177, KL180, KL181 and KL183 were each detected in ≥7 and up to 33 genomes. Notably, these loci were generally associated with multiple STs and geographies; n=14 KL174 genomes with 9 STs from 11 countries, n=8 KL177 genomes with 3 STs (n=6 ST656) from 6 countries, n=7 KL180 genomes with 6 STs from 5 countries, n=9 KL181 genomes with 2 STs from 4 countries, and n=33 KL183 genomes with 22 STs from 14 countries. Further, 9 of the novel loci were detected in genomes sequenced from non-human samples including animal (KL172, KL173, KL175, KL176, KL181, KL183, KL186), environmental (KL178, KL181, KL183) and food sources (KL180, KL183). In particular, 89% of KL181 genomes were sourced from non-human sources (migratory birds, bovine, swine, murine or soil).

While the vast majority of KpSC genomes can be confidently assigned K loci from the existing database, as greater numbers of KpSC genomes become available we anticipate that additional novel K loci will be identified. In particular, it is unlikely that the current database fully captures the diversity present among KpSC circulating in the environment and non-human hosts that remain comparatively underrepresented in current genome collections. Furthermore, *Klebsiella* K loci are thought to undergo frequent reassortment via homologous recombination and IS-mediated horizontal gene transfer both within and outside the *Klebsiella* genus [29, 62], providing a mechanism for the continuous generation of novel loci, and highlighting the need for ongoing population surveillance.

### O locus and O type distributions

In 2018 we implemented KpSC O locus typing in Kaptive, with the capacity to distinguish the O1 and O2 antigen types [30]. In the latest version of Kaptive we have additionally implemented O2 antigen subtype prediction (see **Methods** and **Table 1**). Subsequently, we used Kaptive to explore the distribution of O loci and predicted O types among 12,093 publicly available KpSC genomes (see **Methods** and **Supplementary Table 3**). A total of 11,571 genomes (95.7%) were assigned O locus calls with ‘Good’ confidence or better. Among these, the most common O loci were O1/O2v1 (n=4,015, 34.7%), O1/O2v1 (n=3,413, 29.5%) and O3b (n=1,101, 9.5%). The O1/O2v3 locus, which was added to the database as part of this work (first described in association with O2aeh, GenBank accession MG280710.1 [46]), was identified in only 23 genomes (0.2%).

The distribution of O loci differed by species (**Figure 3**), notably the O1/O2 loci were overrepresented among *K. pneumoniae* compared to other species (*p* < 2.2×10^−16^, OR 64.3, 95% Cl 48.9-86.1, by Fisher’s Exact test), whereas the O3/O3a and O5 loci were underrepresented among *K. pneumoniae* (*p* < 2.2×10^−16^, OR 0.03, 95% Cl 0.025-0.035; *p* < 2.2×10^−16^, OR 0.05, 95% Cl 0.042-0.060; respectively by Fisher’s Exact test). Within *K. pneumoniae*, 47.6% of the isolates carrying an O1/O2 locus were predicted to express an 01 antigen (n=3,521/7,396), while 34.9% (n=2,581) and 16.6% (n=1,224) were predicted to express O2afg and O2a, respectively. No isolates of any species were predicted to express O2aeh and only 75 isolates were predicted to express O2ac (including n=69 *K. pneumoniae*). A single *K. pneumoniae* isolate carried the O1/O2v1 locus in combination with the *wbbY* gene (predicted to result in conversion of O2a to O1), and the *wbmVW* genes (predicted to result in conversion of O2a to O2ac), suggesting that this isolate (NCTC8849, from a respiratory tract specimen collected in the United Kingdom in 1951) may be able to express both the O1 and the O2ac antigens. However, there is evidence that expression of O2ac is thermoregulated, specifically downregulated at 37°C, hence it is likely that this strain expressed the O1 antigen within the human host [46].

**Figure 3:**
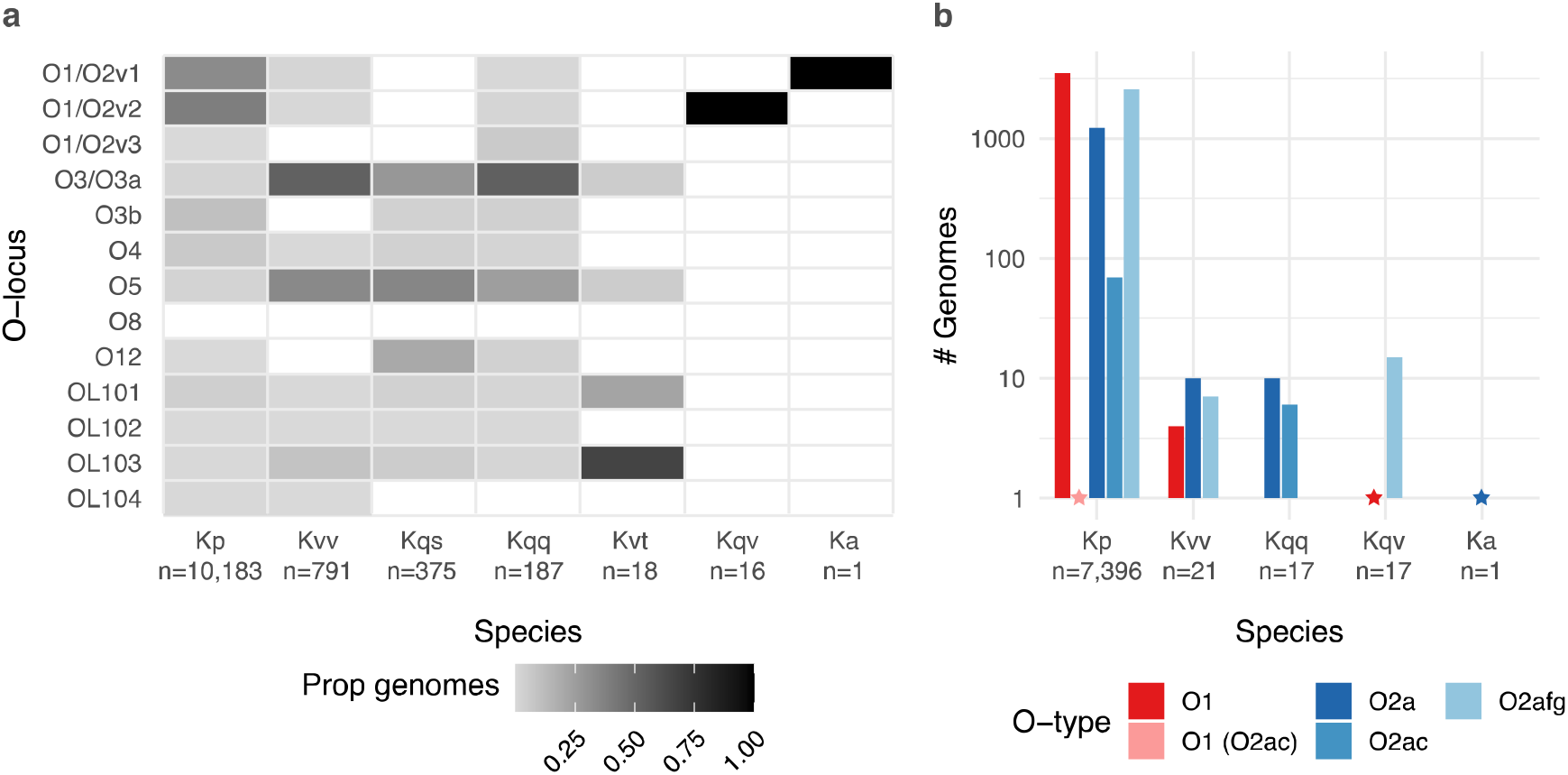
Distribution of O loci and predicted O1/O2 (sub)types by species. **a)** Heatmap showing the proportion of genomes of each species harbouring each distinct O locus. Total sample sizes for each species are indicated below x-axis labels. **b)** Bar graph showing the number of genomes of each species predicted to express O1 and O2 antigens. Only species for which at least one genome was predicted to express an O1 or O2 antigen are shown (total genomes indicated below x-axis labels). Stars indicate the position of bars of size 1. Kp, *K. pneumoniae;* Kvv, *K. variicola* subsp. *variicola;* Kqs, *K. quasipneumoniae* subsp. *similipneumoniae;* Kqq, *K. quasipneumoniae* subsp. *quasipneumoniae;* Kvt, *K. variicola* subsp. *tropica;* Ka, *K. africana*.

Our data also revealed differences in the distribution of O types by isolate source (**Figure 4**). Isolates from human specimens appeared to be enriched for types 01, O2a and/or O2afg compared to those from other hosts and/or environmental sources, which in part likely reflects the dominance *of K. pneumoniae* among clinical specimens. Interestingly, with the exception of liver abscess isolates, the prevalence of O2a was similar across all human-associated source types (8.5% −11.7%) whereas clinical isolates were generally enriched for O2afg compared to gut carriage isolates (22.2% - 30.1% vs. 14.2%, p < 2.2×10^−16^, OR 0.49, 95% Cl 0.42-0.57 using Fisher’s Exact test of gut carriage isolates vs all clinical categories combined) and the prevalence of O1 appeared to be higher among invasive than non-invasive isolates (38.8% among blood/sterile site isolates, 32.0% among respiratory isolates, 28.2% among urinary tract isolates). Most strikingly, the overwhelming majority of liver abscess isolates (84.3%) were predicted to express O1. We speculated that the latter may be driven at least in part by the underlying population structure of *K. pneumoniae* causing liver abscess disease, which is dominated by a small number of so-called ‘hypervirulent’ clones. Indeed our current and previously reported data [28] showed that the most common of these clones (ST23, ST86, ST65 and ST25) were each associated with very high prevalence of O1 (≥81%, **Figure 5**). The data also indicated that the majority of the well-known globally distributed MDR clones and other common clones were each associated with a single dominant O (sub)type, however there was diversity between clones: ST14, ST15, ST20, ST29 and ST101 were each associated with 01 (≥77% each); ST307, ST258 and ST512 were associated with O2afg (≥98%); ST340 and ST437 were associated with 04 (≥91%); ST147 was associated with O2a (68%). In contrast, ST11 (the putative ancestor of ST258 and its descendent, ST512, as well as ST347 and ST437) and the common clones, ST17 and ST37, were each associated with much greater O (sub)type diversity (**Figure 5**).

**Figure 4:**
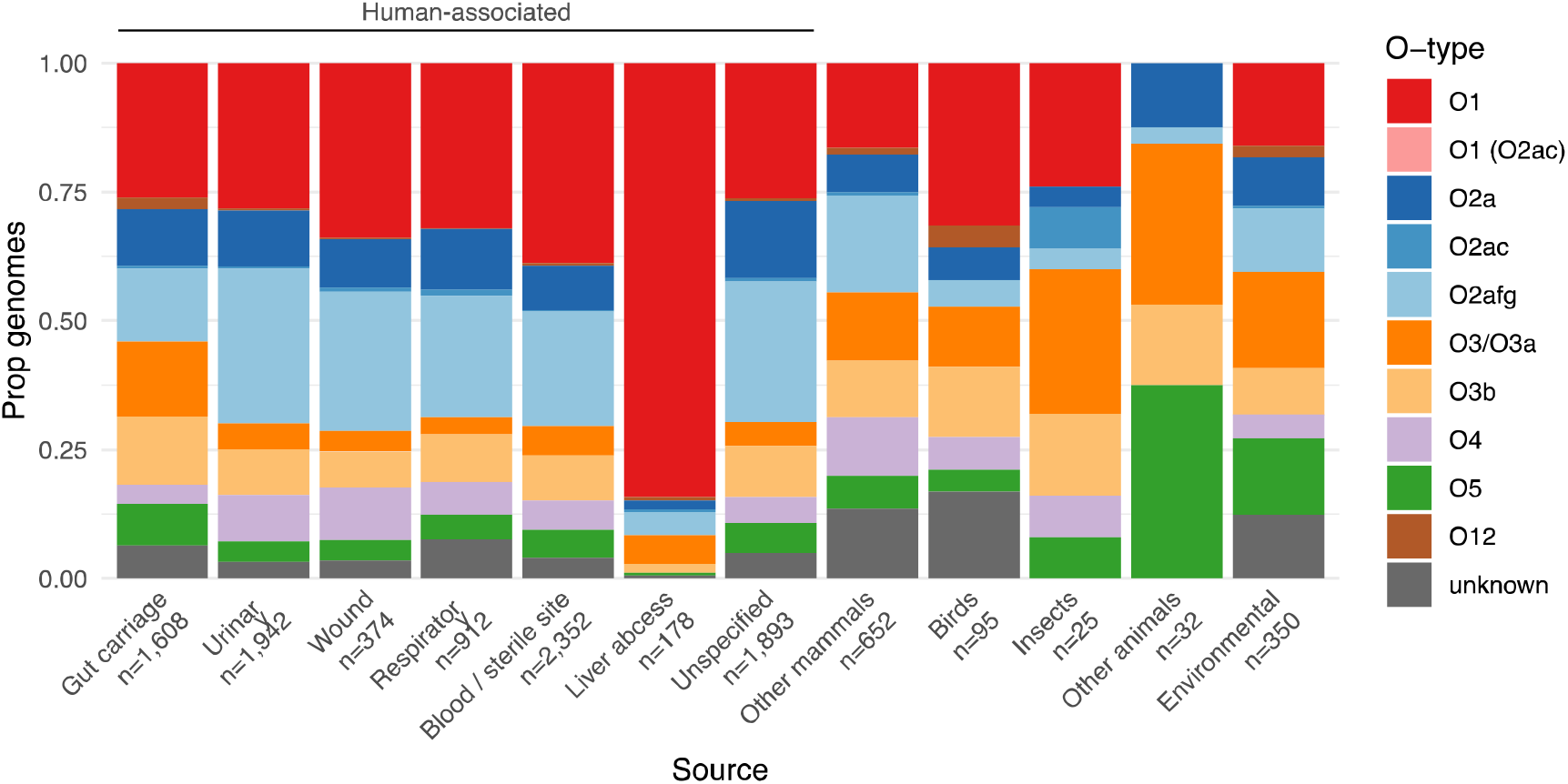
Distribution of predicted O types by isolate source. Bars show the proportion of isolates for each of 12 selected sources of interest that were predicted to encode each O antigen (as indicated in the legend). The total number of isolates representing each source are indicated below the x-axis.

**Figure 5:**
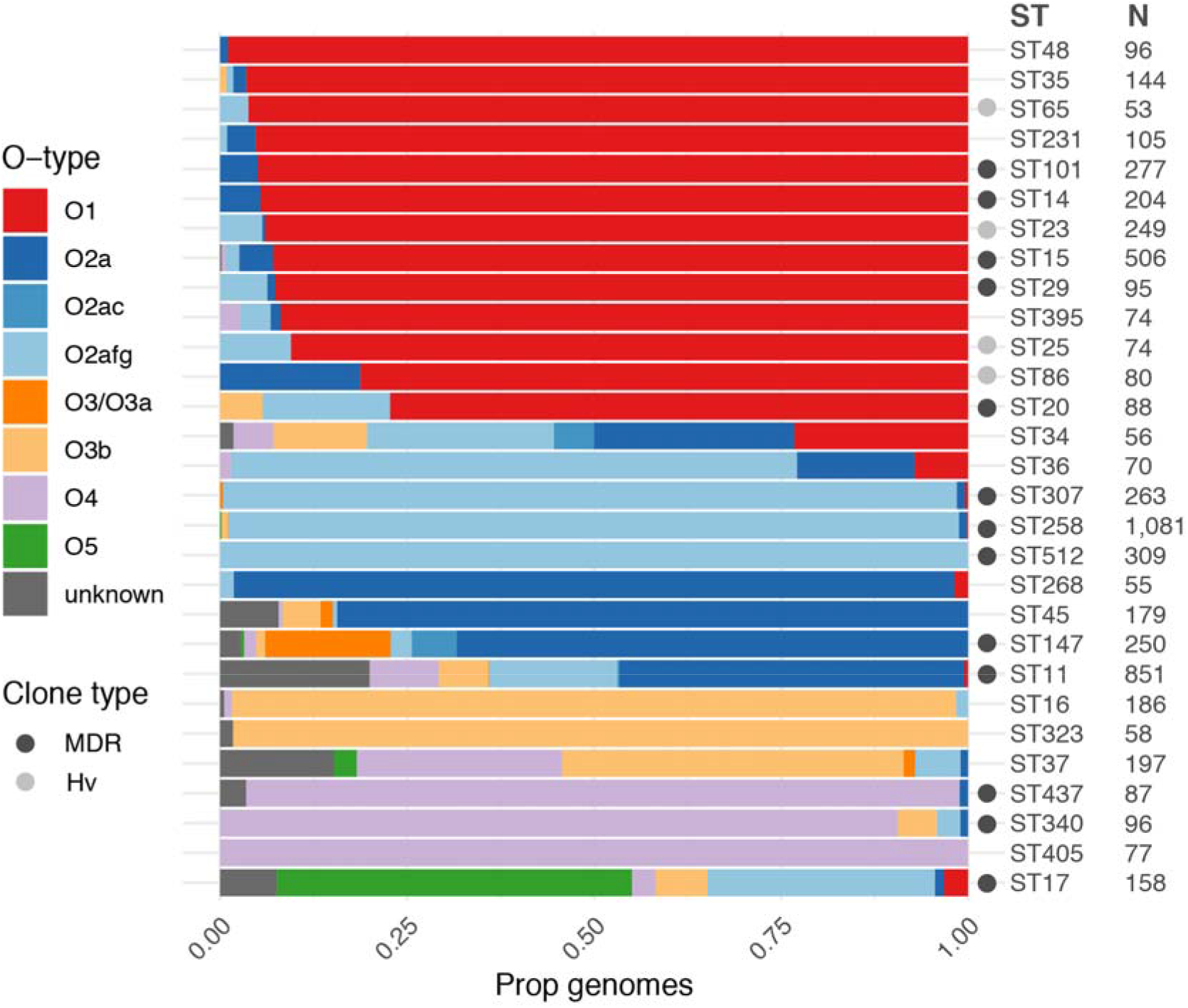
Distribution of predicted O types among common *K. pneumoniae* clones. Bars show the proportion of genomes predicted to express each O (sub)type within each clone (coloured as per legend). Sample sizes are indicated and only clones for which N>50 are shown. Globally distributed multi-drug resistant (MDR) and hypervirulent (Hv) clones are indicated by grey circles as per legend (clones as described previously [1], additionally ST340 and ST437 belong to clonal group 258, ST16 belongs to clonal group 20)).

Previous reports have also indicated a dominance of O1 antigen among human clinical *K. pneumoniae* isolates [27, 28, 63, 64], followed by the 03 group in earlier reports [27, 63] and the O2 group in later reports [28, 64], While few prior studies have distinguished between 02 subtypes, a recent re-analysis of 573 KpSC genome sequences reported in [28] indicated that 42% of the 387 genomes carrying an O1/O2 O locus were predicted to express O2afg, followed by O1, a small number O2a or O2ac, and no O2aeh [46]. Additionally, the O2afg antigen has been previously associated with the ST258, ST512 and ST307 clones [15, 47]. Analysis of 276 neonatal sepsis *K. pneumoniae* genomes from South East Asia and Africa [5] revealed a dominance of 01 (54%) followed by O2afg (14%), 04 (11%), O2a (7%), and less than 5% prevalence of O3/O3a, 03b, 05 and 012. It has been suggested that the lower immunogenicity of O2afg compared to O2a and 01, may provide a selective advantage that has facilitated the widespread dissemination of these clones, i.e. by enabling immune evasion [15,17]; however, we note that our data show that several other highly successful and widespread MDR clones are not associated with O2afg but instead are associated with the highly immunogenic 01 (e.g. ST15, ST20, ST101), indicating that low 0 antigen immunogenicity is not a requirement for widespread dissemination and/or other factors are also playing a role (e.g. the interaction with the polysaccharide capsule).

### Conclusions

*We* present updated KpSC K and O loci databases and an updated version of the Kaptive genotyping tool that allows rapid prediction of O1 and O2 antigen (sub)types from whole genome assemblies. Application of the updated approach to a collection of >12,000 publicly available KpSC genomes indicated key differences in the distribution of O1/O2 antigens by isolate source and strong associations with the underlying KpSC population structure. Further investigation of these trends as well as the broader population sero-epidemiology is warranted to inform the effective design of novel KpSC control strategies. As genomic surveillance of KpSC continues to gain momentum, Kaptive and its accompanying K and O locus databases are poised to play a key role in such investigations.

## Author statements

### Authors and contributors

KEH and KLW were responsible for conceptualisation, supervision, project administration and acquisition of funding. RRW and KLW developed methodology, developed and/or tested software. LMJ performed DNA extractions and isolate sequencing. MMCL and KLW performed investigation, data curation and writing (original draft preparation). All authors performed writing (review and editing).

#### Funding information

This work was supported by the Bill and Melinda Gates Foundation (Investment Grant INV023041 awarded to KEH and KLW). KLW is supported by the National Health and Medical Research Council of Australia (Investigator Grant APP1176192).

