## Supplementary Table 1 for "Kaptive 2.0: updated capsule and LPS locus typing for the *Klebsiella pneumoniae* species complex"

**Supplementary Table 1. Novel K loci identified in this study**

| K locus | Reference genome/read accession | Length (bp) | #CDS | #genomes |
| --- | --- | --- | --- | --- |
| KL171 | GCF_001874715.1 | 24,175 | 21 | 1 |
| KL172 | ERR3449083 | 27,593 | 23 | 1 |
| KL173* | GCF_003990375.1 | 25,719 | 21 | 4 |
| KL174 | ERR315145 | 24,175 | 20 | 15 |
| KL175 | ERR4367681 | 28,533 | 25 | 1 |
| KL176* | ERR3448903 | 25,166 | 19 | 5 |
| KL177 | T7-221 | 29,929 | 25 | 8 |
| KL178 | T7-392 | 28,865 | 24 | 3 |
| KL179 | GCF_900407305.1 | 29,218 | 23 | 2 |
| KL180 | T7-177 | 30,215 | 24 | 7 |
| KL181* | ERR4367597 | 27,966 | 23 | 9 |
| KL182 | JAJHNT000000000 | 26,104 | 22 | 2 |
| KL183 | T7-391 | 22,626 | 19 | 33 |
| KL184* | ERR4367641 | 26,378 | 24 | 2 |
| KL185* | GCF_900493845.1 | 26,462 | 21 | 3 |
| KL186 | GCF_002247665.1 | 25,893 | 21 | 5 |

*K loci where IS were manually removed from the reference sequence
